# Diet links gut chemistry with cancer risk in C57Bl/6 mice and human colorectal cancer patients

**DOI:** 10.1101/2025.01.27.635083

**Authors:** Ziv Cohen, Jiahn Choi, Karina Peregrina, Saad Khan, Sarah Wolfson, Cherrie Sherman, Leonard Augenlicht, Libusha Kelly

## Abstract

The gastrointestinal tract is a complex ecosystem in which host tissues, microbial communities, and dietary inputs interact to shape metabolic outputs and epithelial homeostasis. Western-style diets, characterized by high fat and protein and low micronutrient content, represent a sustained ecological perturbation linked to colorectal cancer (**CRC**), yet specific mechanisms that impact risk remain poorly defined. Here, using a purified Western-style diet (**NWD1**) that induces sporadic intestinal and colon tumors in wild-type C57BL/6 mice we demonstrate how chronic dietary disturbance restructures gut community composition and sulfur metabolism, and how these changes modulate intestinal stem cell responses. Mice fed NWD1 for 24 weeks exhibited consistent shifts in ecosystem function, including a threefold increase in fecal sulfide production (P < 0.00001) and expansion of *Erysipelotrichaceae* family taxa. This altered chemical landscape associates with increased expression of mitochondrial sulfide oxidation pathways in Lgr5^hi^ intestinal stem cells. Meta-analysis of human CRC cohorts revealed concordant enrichment of *Erysipelotrichaceae*, alongside established CRC-associated taxa such as *Solobacterium moorei*, indicating conserved ecological signatures across systems. Together, these findings support a model in which the Western-style diet drives a persistent shift in gut ecosystem structure and function toward a sulfide producing state that challenges epithelial homeostasis, with conserved microbial and metabolic configurations emerging prior to overt disease in mouse models.

## Introduction

In humans, a Western diet is broadly characterized by higher intake of fat, animal protein, and refined grains and sweets; and lower intake of fiber, fruits, vegetables, vitamin D, calcium, and methyl-donor nutrients^1,2^. Western-style dietary patterns promote a pro-tumorigenic environment, and in concert with the microbiome, can induce inflammation, changes in host epithelial signaling pathways, immune modulation, and production of metabolites that increase the risk of developing colorectal cancer^3–5^. One such metabolite is hydrogen sulfide. Increased sulfide production is linked to Inflammatory Bowel Disease (IBD), inflammation, and colon tumor development^6–8^, and at high concentrations is toxic to colonocytes^9,10^.

A consequence of perturbations to the gut environment by Western-style diets and microbiota is the development of sporadic cases of colorectal cancer (CRC) ^11^. Most CRC cases are sporadic and influenced by modifiable environmental factors ^11^. The gut microbiome is associated with key drivers of tumorigenesis, including inflammation, changes in host epithelial signaling pathways, immune modulation, and production of metabolites that increase the risk of developing CRC^3–5^. Numerous studies have reported microbes associated with early and late-stage CRC^12–14^. Western-style dietary patterns promote CRC development, progression, and mortality in both human and mouse studies and can shape microbiome composition and function to promote or exacerbate disease^1–3,15–17^. Furthermore, adherence to a “sulfur microbial diet”, characterized by increased intake of processed meats and decreased intake of fruits and vegetables, is associated with sulfide-producing bacterial species and increased incidence of premalignant adenomas^18^. Thus, global changes to diet, and therefore sulfide, can be detrimental to the gut ecosystem, changing microbiome composition as well as the human gut lumen itself.

Many different rodent diets are used to examine the influences of Western-style diets on intestinal homeostasis and disease; here, we focus on a Western-style purified diet (NWD1) in which multiple dietary components are adjusted to expose mice to levels of each nutrient that are epidemiologically linked to those consumed by individuals at high risk for CRC in both developed and developing countries^19,20^. Relative to the standard purified mouse diet AIN76A, NWD1 has lower levels of calcium, vitamin D, fiber, and methyl-donor nutrients, and increased fat content that reflect levels linked to increased CRC risk associated with Western human diets. Notably, while mice fed NWD1 gain some weight, they do not become obese on this diet, as obesity has been shown to be an independent risk factor for colorectal cancer^21,22^. Feeding NWD1 accelerates and amplifies intestinal tumors in many genetic models^23–26^ and is unique in reproducibly causing sporadic intestinal tumors in wild-type mice in the absence of genetic predisposition or carcinogen exposure. These mice rarely, if ever, develop such tumors when fed other diets over their lifespan^20,26^. Approximately 20% of wild type mice fed NWD1 develop intestinal tumors^2^, reflecting the etiology, incidence, frequency, and lag-time with developmental age of sporadic human colon tumors, which are the most prevalent form of the disease and the form most influenced by long-term dietary patterns^27,28^.

Part of the mechanism by which NWD1 reshapes the gut luminal environment is through the alteration of gene expression in intestinal stem cell (ISC) populations. Lgr5^hi^ ISCs at the crypt base are canonical stem cells that give rise to all lineages, maintaining mucosal structure and function^29,30^. NWD1 reprograms these stem cells, including repressing mitochondrial oxidative phosphorylation and tricarboxylic acid cycle pathways. This represses their stem cell functions, slows developmental maturation of the daughter cells of these Lgr5^hi^ ISCs, and gives rise to lineages that are remodeled to be proinflammatory and protumorigenic^28,31,32^. The NWD1 diet thus allows the examination of complex dietary changes as a targeted environmental risk factor on the intestinal epithelium and microbiome.

Here, we examine mechanisms underlying increased CRC risk by treating the Western diet as a sustained ecological perturbation to the gut ecosystem. We used a murine model fed NWD1 to investigate dietary effects on the microbiome, sulfide production, and mRNA transcription in intestinal epithelial stem cells, and followed this with a nine-study meta-analysis of human metagenomes from human CRC cases (n=956) and controls (n=694)^33–42^. Mice fed NWD1 experienced alterations in the composition of their microbiomes, with higher relative abundances of *Erysipelotrichaceae*, which in human gut microbiomes includes bacteria associated with CRC^18,43,44^. Despite differences in microbiome composition among male and female mice, mice fed NWD1 produced more sulfide than mice fed AIN76A and had higher expression of mRNA in the mitochondrial sulfide oxidation pathway. Finally, changes in the microbiome composition in the mice, which had not developed tumors by the end of the study, mimicked the changes in metagenomes from human CRC patients. Together, these findings demonstrate that sulfide production is an ecosystem-level readout of dietary exposure and suggest that shifts in microbial community function driven by diet and chemistry precede disease and may thus provide targets for risk stratification or intervention.

## METHODS

### Animals and Fecal Sample Collection

Inbred C57BL/6 mice were housed in a barrier facility at Albert Einstein College of Medicine, with all experiments conducted under protocol 00001045 of the Einstein Institutional Animal Care and Use Committee (IACUC). Twenty C57BL/6 mice were randomized into four cohorts: five female and five male mice were fed the AIN76A diet, and five female and five male mice were fed the NWD1 diet from weaning. Each cohort of mice (female AIN, male AIN, female NWD1, male NWD1) was housed in its own cage, with a total of five mice per cage. The mice had *ad libitum* access to their respective diets and water. Fresh fecal samples were collected from each mouse at 2, 4, 8, 12, 16, 20, and 24 weeks, respectively, to assess changes in microbial composition over time.

### Dietary Composition

Detailed dietary composition information is in Newmark *et al* 2001^2^. Briefly, nutrients are in units, weight, or percent of diet by weight for each component per gram of diet (**Table 1**). Numbers in parentheses are based on nutrient densities from the average human intake on a 2000 Calorie (kcal) diet. NWD1 values are based on either lower quartile nutrient intake or average human intake on a 2000 Calorie (kcal) diet. Two exceptions are fat (in percent calories in the parentheses), and fiber (“grams per day” in the human diet). Methionine and cysteine are supplied by the exclusive protein source, casein, with excess methionine and cysteine relative to baseline listed in the table.

**Table 1.** Dietary Composition.

| Ingredients, % (wt)<br>or amount (wt) | Control<br>AIN76A | NWD1 |
| --- | --- | --- |
| Fat (corn oil), % | 5 (13) | 20 (40) |
| Calcium, mg/g | 5 (2700) | 0.5 (220) |
| Vitamin D<sub>3</sub>, IU/g | 1 (600) | 0.11 (50) |
| Phosphorus (PO<sub>4</sub>), mg/g | 4 (2200) | 3.6 (1600) |
| Fiber (cellulose), % | 5 (25) | 2 (9) |
| Folic acid, µg/g | 2 (1100) | 0.23 (100) |
| dl methionine, % | 0.3 | -- |
| l cysteine, % | -- | 0.3 |
| Choline bitartrate, % | 0.2 | 0.12 |
| kcal/g (approximate) | 3.6 | 4.5 |

### Fecal Sulfide Measurement^45,46^

Freshly voided fecal pellets were collected from an additional cohort of seven-month-old mice fed AIN (n=3 female, n=5 male) or NWD1 (n=3 female, n=5 male) on three separate days within a one-week period. The pellets were immediately submerged in 157 µL of 28 g/L zinc acetate. Samples were then diluted with 1.26 mL water and reacted with 113 µL of Cline’s reagent (N,N-dimethyl-p-phenylenediamine and FeCl_3_). The samples were incubated for 20 minutes at room temperature for color development and then centrifuged to remove particulates. Next, 100 µL of the supernatant was dispensed into Costar clear flat-bottomed 96-well plates (Corning). Absorbance was recorded at 670 nm, and the H_2_S concentration was quantified by UV–Vis detection on a BioTek Synergy H4 plate reader (BioTek Instruments) using Gen5 version 1.11 software.

## Data Availability

Microbiome sequences and metadata are deposited in the Sequence Read Archive under PRJNA981173 (https://www.ncbi.nlm.nih.gov/bioproject/PRJNA981173). The RNA sequencing data is available in the Gene Expression Omnibus under Accession GSE186811 (https://www.ncbi.nlm.nih.gov/geo/query/acc.cgi?acc=GSE186811).

## Sequencing

Freshly voided fecal pellets were collected from individual mice every two weeks. Bacterial DNA was extracted from the fecal pellets using the Qiagen PowerFecal kit (Germantown, Maryland, USA). Fecal pellet microbiome 16S V4-V5 sequencing was carried out by the Integrated Microbiome Resource (Halifax, Canada).

## Bioinformatics

### Taxonomic Community Profiling with Metadata - QIIME

Microbiome analyses were performed with QIIME 2 q2cli version 2022.8.0^47^; no original code was generated for the project. Sequences were demultiplexed using the q2-demux plugin^48,49^. The sequences were denoised and filtered with q2-deblur denoise-16S using a –p-trim-length of 301^50–52^, and a feature table was created^53–55^. The sequences were aligned with mafft using the align-to-tree-mafft-fasttree pipeline from the q2-phylogeny plugin^56–58^. Diversity metrics (Faith’s phylogeny index, evenness) were calculated using the q2-diversity plugin with a –p-sampling-depth of 5350 to ensure all samples were included^51,59–62^. Taxonomic analysis at the phylum, family, and genus levels was carried out with the q2-feature classifier trained on the Greengenes 13_8 99% OTUs reference sequences^63–66^.

### Assessing Microbial Community Relationships to Metadata - MaAsLin2

Microbial community features were assessed for relation to metadata using MaAsLin2 run in R version 4.2.2 (2022-10-31 ucrt)^67^. Fixed effects were “Diet” and “Sex”; min_prevalence = 0.1, and min_abundance = 0.0001 were used.

### Bulk RNAseq Analysis of Lgr5^hi^ Intestinal Stem Cells

Mice that were Lgr5^cre:ER-Egfp^ were fed AIN76A continuously for three months or switched to the NWD1 diet for the final four days as reported^28^. At the time of euthanasia, excised intestines were opened longitudinally and rinsed with cold saline, crypts were isolated, and single-cell suspensions were prepared and sorted by FACS to collect the 2% of cells with the highest expression of Egfp marker, which are intestinal crypt base stem cells^29,30^. Bulk RNAseq was done with the RNA isolated from these cells and reported^28^. The data are available in the Sequence Read Archive under PRJNA981173.

### Human CRC Case-Control Meta-Analysis

We retrieved data on colorectal cancer gut microbiomes with matched healthy controls from nine different studies ^33–42^ using a curated library^68^. These samples were uniformly preprocessed and taxonomically annotated using MetaPhlAn3^69^, leading to a total of 956 colorectal cancer and 694 healthy control microbiomes. Briefly, we conducted a meta-analysis by first calculating Mann-Whitney association area under the curve (AUC) statistics for each taxon with each study. The AUC gives a measure of the degree and direction of association of that taxon for health (AUC>0.5) or disease (AUC<0.5). For each taxon, we then calculated a weighted average of the taxon’s AUC score across each dataset (weighted by the inverse square root of the sample size of the study). We then conducted single-sample t-tests for each taxon to test the null hypothesis that the mean AUC score is equal to 0.5 (no association). We reported those taxa for which the null hypothesis was rejected at alpha = 0.05 as being consistently linked to CRC (or healthy). The statistical basis for the method is described in **Supplementary Methods**.

### Analysis of Sulfide-Producing Genes

To identify the presence or absence of sulfidogenic genes in *Erysipelotrichaceae* taxa, databases of homologous sequences for CBS (K01697), CSE (K01758), cysK (K01738), cysM (K12339), cyuA (COG3681), dsrA (K11180), dsrB (K11181), malY (K14155), metC (K01760), sseA (K01011) and tnaA (K01667) were constructed and searched against each reference genome. A reference protein database for each gene was created as reported^46^. Briefly, for each gene, 500 closely related bacterial orthologues were aligned using MUSCLE with default parameters (MUSCLE 3.8.1551)^70^. Alignments were used to construct hidden Markov models using hmmbuild with default parameters (hmmer 3.3.2)^71^ and searched against the genome using hmmsearch with a stringent cutoff of 1□×□10−10.

## Results

### Temporal differences among mouse microbiomes induced by diet

To determine compositional differences among the microbiomes of the mice fed the NWD1 (Western) and AIN (control) diets, we represented taxonomy at the bacterial family level. In week 2, there was a higher relative frequency of *Bacteroidaceae* in mice fed the AIN diet, contrasted with higher levels of *Erysipelotrichaceae* in mice fed NWD1 (**Fig. 1a**). By week 12 (**Fig. 1b**), there was a decrease in the relative frequency of *Bacteroidaceae* and *Muribaculaceae* (previously named S24-7)^72^ in the AIN-fed mice, with an increase in the relative frequency of *Erysipelotrichaceae* and *Ruminococcaceae.* In the NWD1-fed mice, the opposite was observed: increased relative frequency of *Muribaculaceae,* with some mice exhibiting increased *Bacteroidaceae* or *Erysipelotrichaceae.* By week 24 (**Fig. 1c**), some of the lower relative frequency families increased in the AIN mice. Additionally, the *Verrucomicrobiaceae* and *Ruminococcaceae* decreased in relative frequency. In the NWD1 mice, *Erysipelotrichaceae, Bacteroidaceae,* and *Muribaculaceae* maintained high relative frequencies in the community. Across all weeks, we observed higher alpha diversity in the AIN mice (Faith’s phylogenetic diversity), as well as higher community evenness (**Supplementary Figures 1-3**).

**Figure 1.**
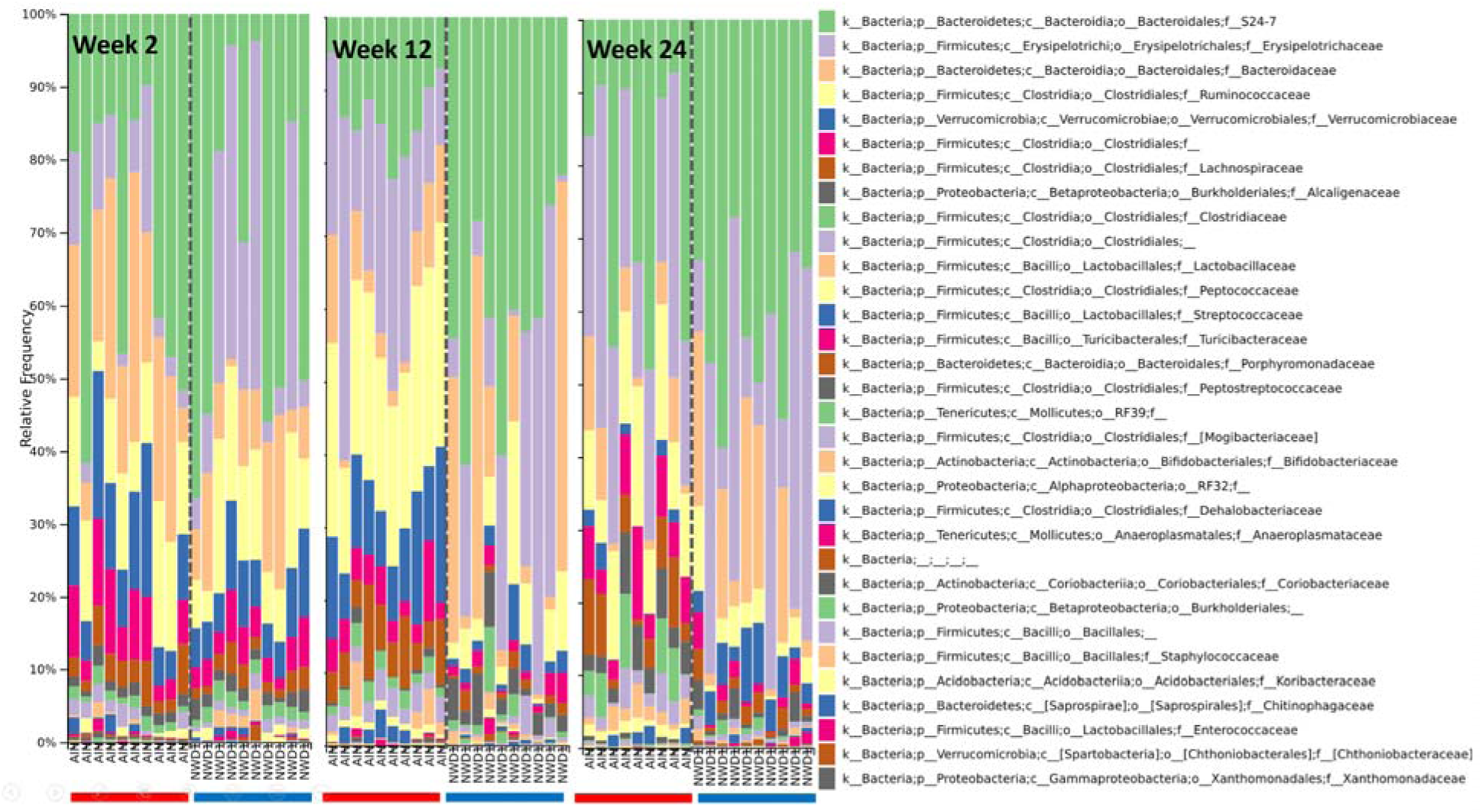
Composition of microbiomes at the bacterial family level from male and female mice fed AIN76A or NWD1 diet at week 2, 12, and 24. Columns represent the taxonomy of mice microbiomes separated by diet at week 2, week 12, and week 24 respectively. The microbiome samples from mice fed NWD1 show greater levels of the *Erysipelotrichaceae* family, shown in lavender.

### Higher relative abundance of *Erysipelotrichaceae* in NWD1-fed mice compared to AIN

We plotted the relative abundance of the most abundant bacterial families (**Supplementary Fig. 4, 5**) in the mice fed AIN and NWD1, finding that the relative abundance of *Erysipelotrichaceae* members (**Figure 2**) was higher in mice fed NWD1 (MaAsLin2, *P = 0.019917, Q = 0.025892*).

**Figure 2.**
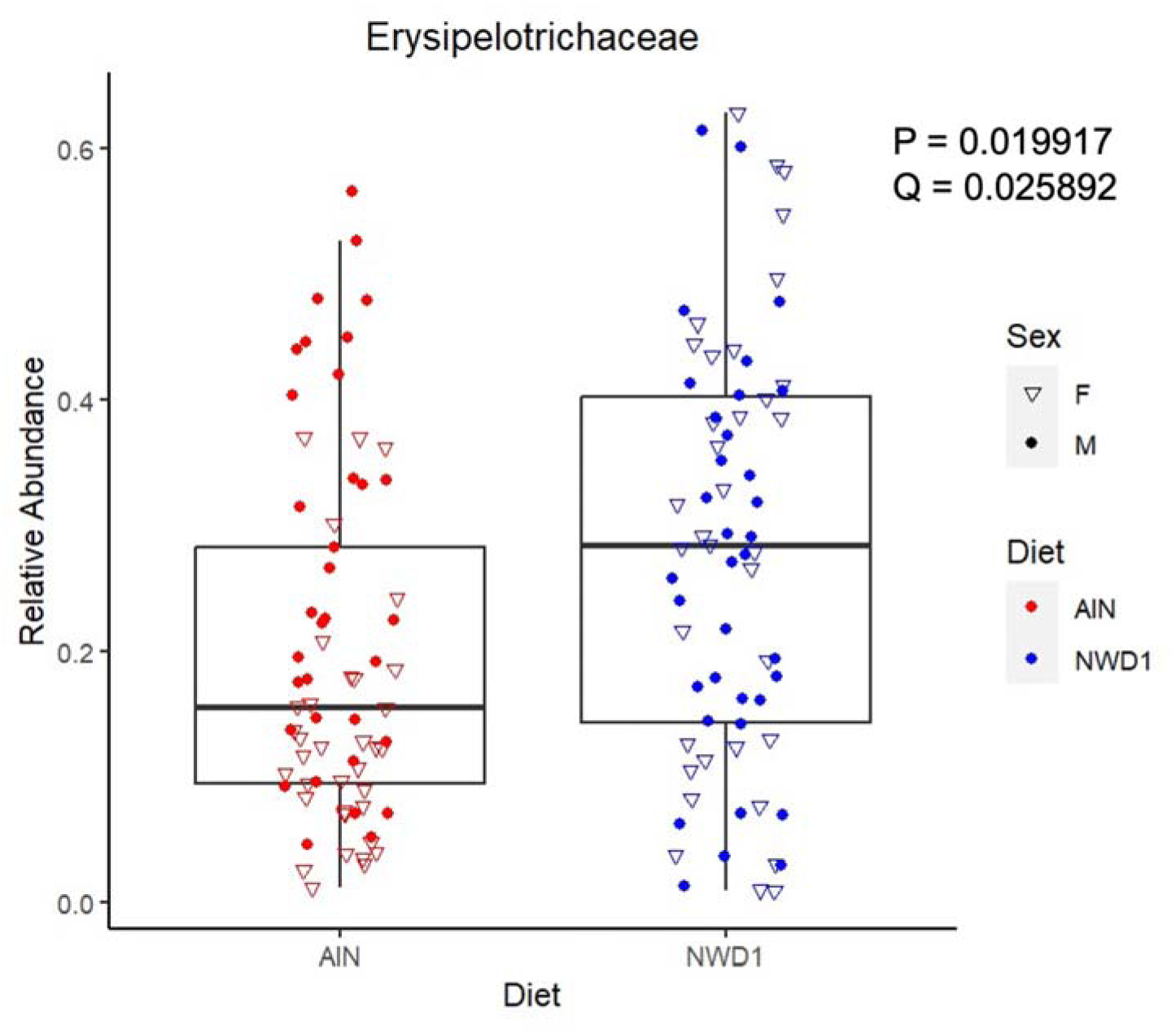
The relative abundance of the *Erysipelotrichaceae* family is higher in mice fed NWD1 vs. AIN. Each point in the plot represents a sample taken over 24 weeks. Samples from mice on the AIN diet are indicated in red, NWD1 samples are indicated in blue. Samples from male mice are represented by open triangles, female mice by filled dots. *Erysipelotrichaceae* relative abundance is significantly higher in mice fed NWD1 (MaAsLin2 default *FDR < 0.25, P = 0.019917, Q = 0.025892*).

### Sulfide production varies by diet but not by sex

Recent reports demonstrate that diet alters sulfide levels, with potential links among microbiome function, dietary modification, and pathophysiological changes^9,73–75^. Because elevated bacterial hydrogen sulfide production is linked to both dietary pattern and colorectal cancer^76–78^, we determined whether NWD1 alters sulfide production by assaying fecal sulfide of male and female mice fed the AIN or NWD1 diet. Mice fed the NWD1 diet had an approximately threefold, highly significant increase in fecal sulfide compared to mice fed the AIN control, regardless of sex (**Fig. 3**, Wilcoxon Rank Sum Test, *P<0.00001*). There was no difference in sulfide production among mice of different sexes fed the same diet.

**Figure 3.**
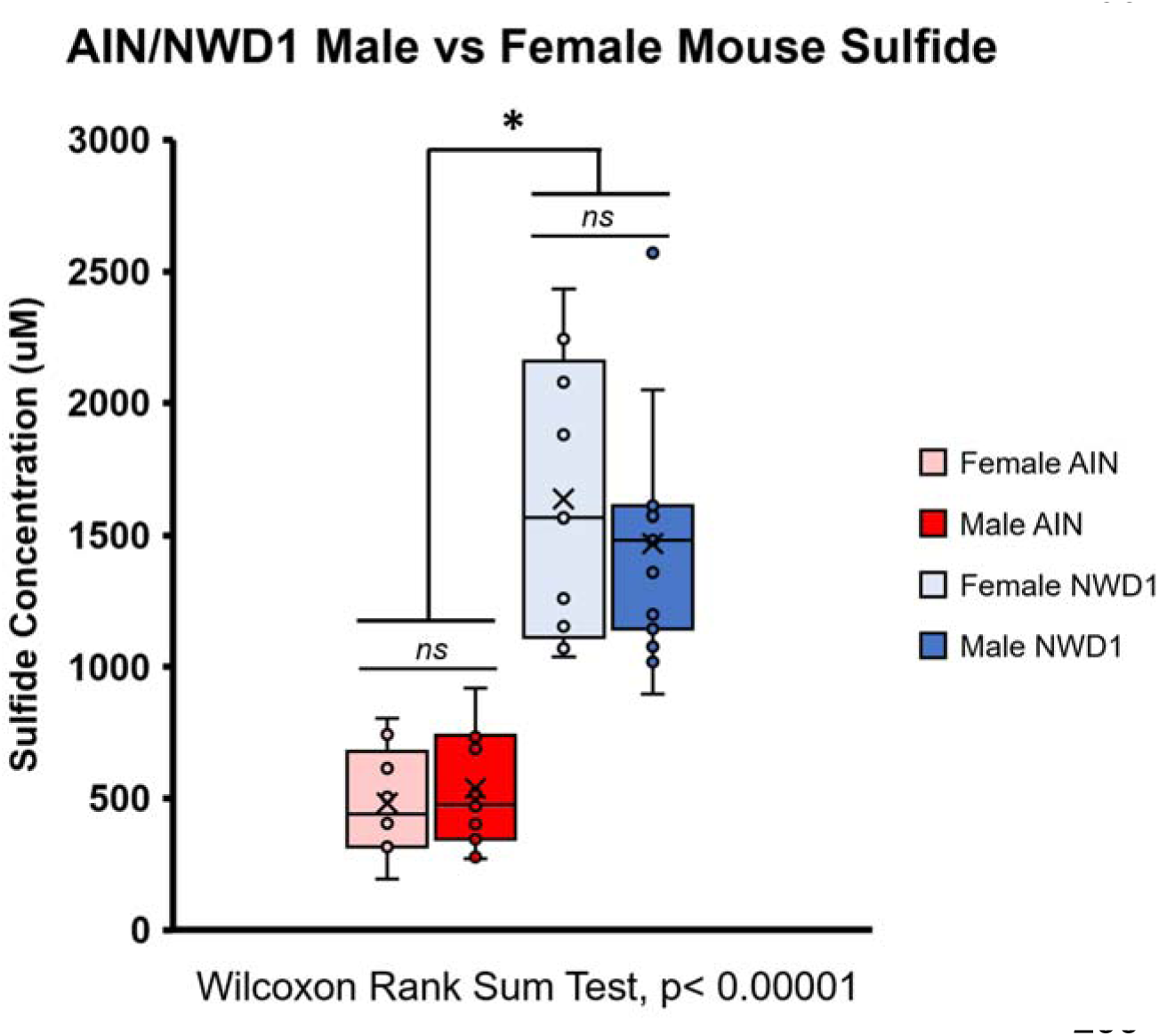
Fecal sulfide is higher in mice fed NWD1. Seven-month-old male and female mice were fed either the control AIN76A diet or NWD1, a Western-style, colorectal cancer-promoting diet from weaning. A total of three female mice were fed AIN, three female mice were fed NWD1, five male mice were fed AIN, and five male mice were fed NWD1. Fresh fecal pellets were collected on three separate days during a one-week period and their fecal sulfide was assessed via a modified Cline assay. Fecal sulfide was significantly higher in the mice fed the NWD1 diet than in those fed the AIN diet (Wilcoxon Rank Sum Test, *P< 0.00001*), but there was no difference between male and female mice on the same diet for both AIN and NWD1.

### Lgr5^hi^ Intestinal Epithelial Stem Cells Undergo Adaptive Changes to Elevated Sulfide

We previously reported that intestinal stem cells and lineages undergo substantial and dynamic transcriptional reprogramming in response to NWD1. This includes adaptive responses to altered nutrients in the diet (e.g. higher fat and lower calcium, epigenetically upregulating genes involved in fat metabolism and calcium uptake). The reprogramming of Lgr5^hi^ ISCs also altered their function as stem cells, recapitulated by targeted knockout in these cells of a key gene downregulated by NWD1, triggering a cascade of further remodeling of progenitor cells and lineages^28^. To determine if the stem cells adapt to the increased sulfide produced in response to NWD1, we interrogated bulk RNAseq data of Lgr5^hi^ intestinal stem cells from AIN-and NWD1-fed mice for the expression of genes important in sulfide metabolism, namely, the mitochondrial sulfide oxidation pathway^28^ (**Fig 4a**). After four days of shift to NWD1, there was no significant change in the expression of sulfide-quinone reductase-like protein (Sqrdl), which encodes the transporter of sulfide into the mitochondria. However, three genes necessary for sulfide metabolism and its detoxification by oxidation to sulfate, ethylmalonic encephalopathy 1 (Ethe1), thiosulfate transferase (TST/rhodanese), and sulfite oxidase (Suox), were upregulated by NWD1 (**Fig 4b**).

**Figure 4.**
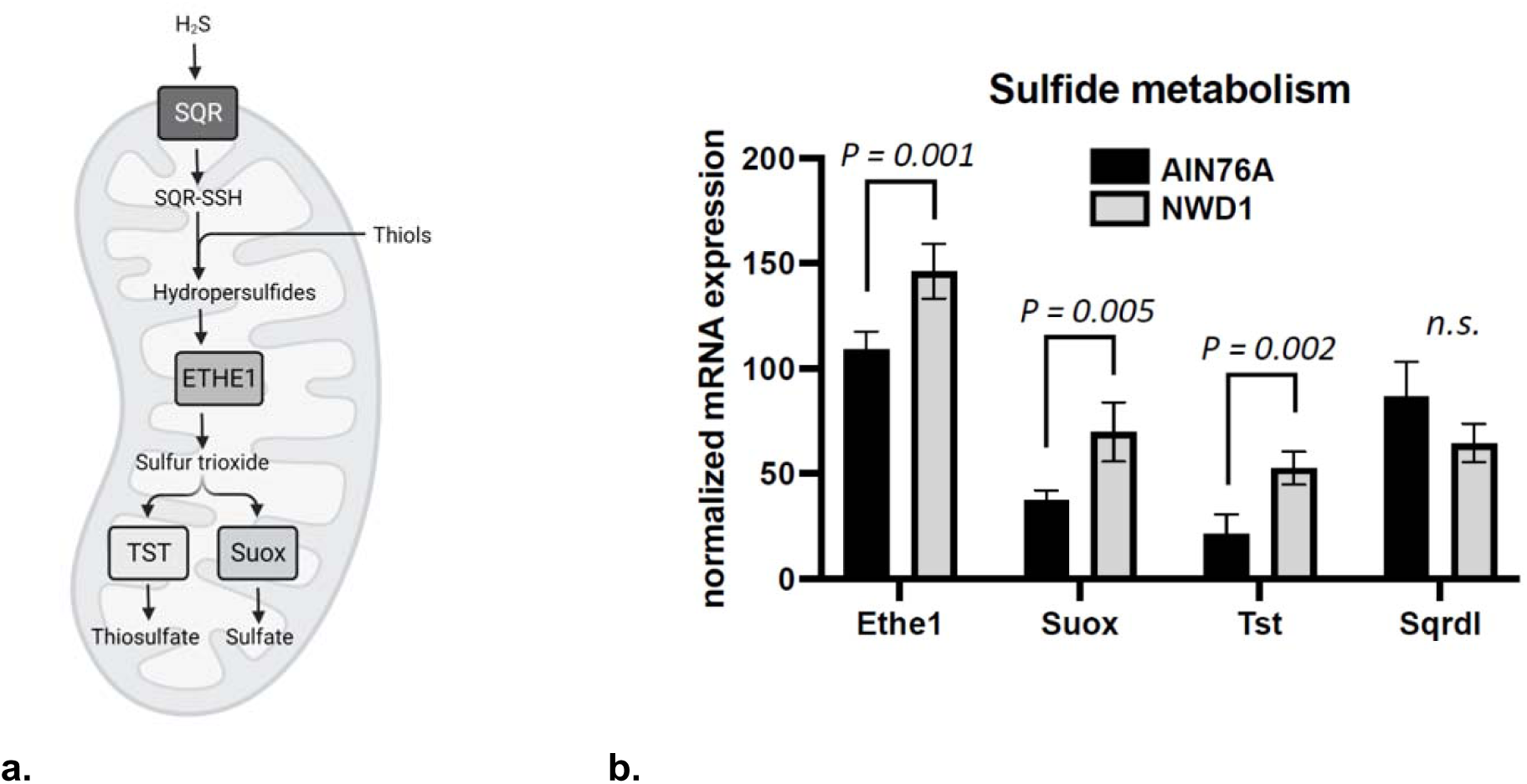
Bulk RNAseq analysis of Lgr5^hi^ intestinal stem cells from AIN- or NWD1-fed mice shows alterations in expression of the mitochondrial sulfide oxidation pathway. a) Model of mitochondrial sulfide oxidation pathway with key enzymes (boxes) and sulfur species (plain text) labeled. Modified from Kolluru et al 2023. b) Three enzymes involved in mitochondrial oxidation, Ethe1, Suox, and Tst, were elevated in Lgr5^hi^ stem cells from Lgr5^cre:ER-Egfp^ mice fed NWD1, while there was no significant difference in Sqrdl.

### *Erysipelotrichaceae* species are associated with disease in CRC studies

Based on the finding that *Erysipelotrichaceae* taxa were associated with the NWD1 diet, we next determined whether *Erysipelotrichaceae* taxa were similarly associated with human colorectal cancer patients. Using 1,650 metagenomes from nine studies spanning multiple countries of CRC patients vs. controls, we determined the associations (average AUC by study) of ten species of the *Erysipelotrichaceae* family to CRC or health. Nine of ten (9/10, 90%) species trended towards being disease associated (**Fig. 5a),** which includes four species that are statistically significantly associated with disease. Each species had multiple cysteine desulfhydrase genes, which have the capacity to produce sulfide (**Fig. 5b).**

**Figure 5.**
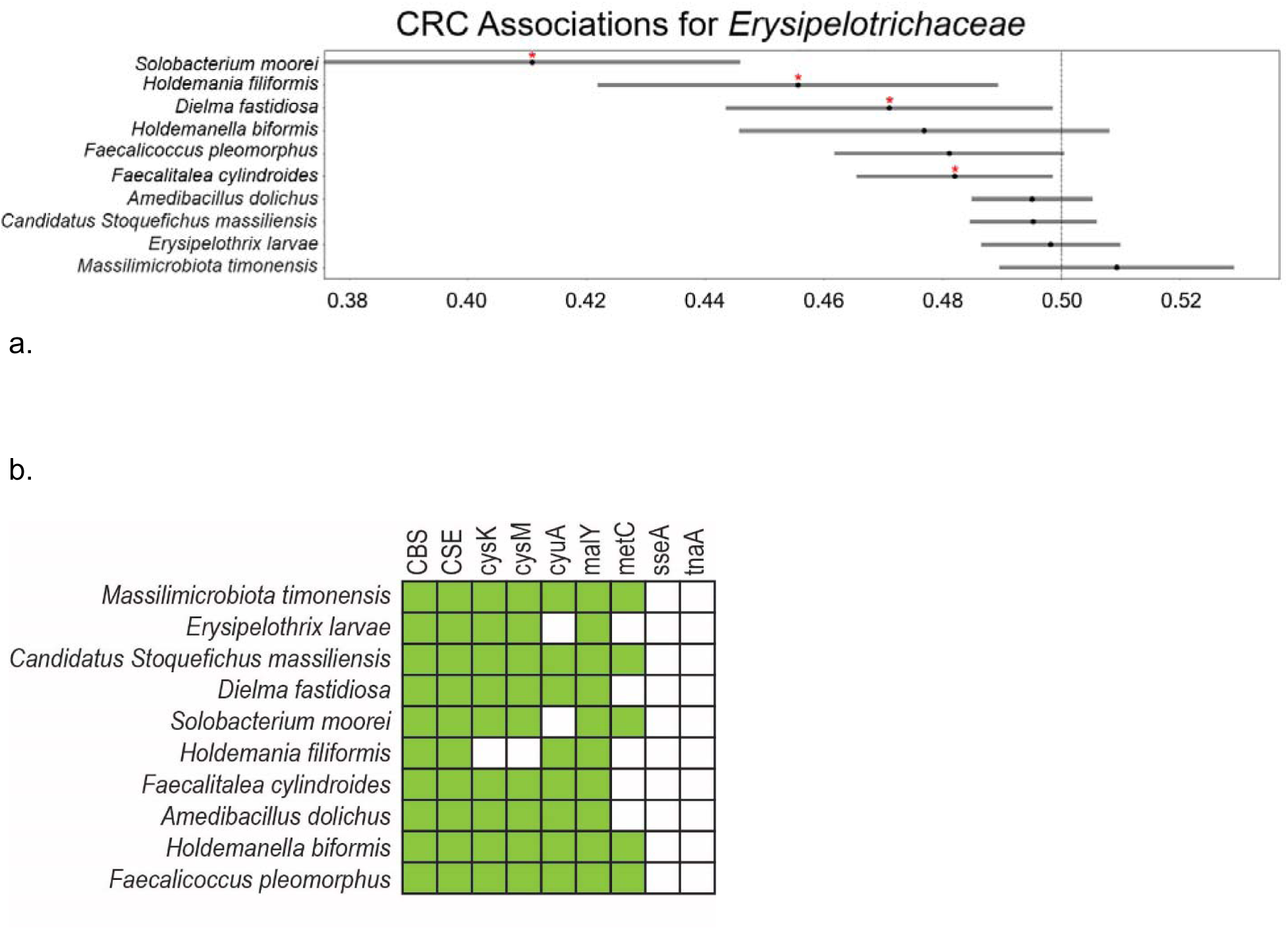
*Erysipelotrichaceae* associations with health and disease in colorectal cancer. a) Nine case-control CRC microbiome studies were compiled and *Erysipelotrichaceae* associations with control or CRC were evaluated using Mann-Whitney AUC scores. Cross-study AUC scores are plotted for all identified *Erysipelotrichaceae* family members. *Erysipelotrichaceae* as a whole trend towards disease association, with mean scores consistently below 0.5. Scores statistically lower than 0.5 that do not cross the midline (**\***) indicate species that are significantly CRC associated (Student’s t-test, *P<0.05*). b) *Erysipelotrichaceae* taxa were assessed for presence or absence of sulfidogenic cysteine desulfhydrase genes. Databases of homologous sequences for cysteine desulfhydrase genes were created and searched against each reference genome with a stringent cutoff of 1□×□10^−10^. Filled boxes indicate the presence of at least one copy of a gene in the genome.

## DISCUSSION

### Diet changes the gut ecosystem to make it more prone to neoplasm

Dietary composition is an environmental perturbation that can either promote or inhibit pathological changes via alterations in the composition of gut microbes and their metabolism of dietary nutrients. Pathological microbe/diet interactions in the host ecosystem can send signals to the mucosa that, over time, can create a cycle of host tissue remodeling that promotes inflammation and alters epithelial homeostasis in the absence of carcinogen exposure or genetic predisposition to tumor development. Clear epidemiological data establishes the importance of dietary patterns as a major determinant of sporadic colorectal cancer incidence in human populations. Here, we demonstrate that feeding C57BL/6 mice the Western-style, colorectal cancer-promoting NWD1 diet reshapes multiple aspects of ecosystem functioning including increases in the relative abundance of *Erysipelotrichaceae*, higher fecal sulfide production, and the adaptive intestinal stem cell response to increased sulfide. We demonstrated that *Erysipelotrichaceae* have a higher relative abundance in NWD1-fed mice, which had not yet developed tumors, and were disease-associated in human CRC metagenomes. Since CRC development is associated with Western-style diets that are higher in sulfur-containing substrates, future work will determine the relationship between *Erysipelotrichaceae,* sulfide production from dietary sources, and CRC risk and development in humans.

### Dietary modification results in taxonomic and chemical changes in the gut microbiome

Microbial diversity was reduced in the mice fed the Western-style diet (Supp. Fig. 1-3). When bacterial composition was determined in the mice on the Western-style (NWD1) and control (AIN) diets at the family level, some mice harbored upwards of 60% relative abundance of *Erysipelotrichaceae. Erysipelotrichaceae* are highly immunogenic, disease-linked, and enriched in colorectal cancer, as well as strongly associated with the consumption of a higher fat diet^18,43,79–81^. Our study further corroborates this, as the NWD1 diet contains a higher proportion of calories from fat, although not as high as in commonly used, very high (60%) fat diets that produce mouse obesity^82^.

Importantly, we show that the *Erysipelotrichaceae* family is substantially increased in relative abundance by NWD1, which coincides with the elevation of fecal sulfide levels. We demonstrated for the first time that NWD1 is a sulfidogenic diet: mice fed NWD1 had a threefold higher level of fecal sulfide than mice fed AIN76A. These results suggest that diet is the strongest determinant of gut sulfide production, as sulfide concentrations were significantly higher in NWD1-fed mice irrespective of individual variation in microbiome composition. Microbial sulfidogenesis has been implicated in the development of colorectal cancer precursors, demonstrating that microbiota and their metabolism integrate environmental and dietary risk factors^83^. Thus, we posit that sulfide production can be considered a community-level functional readout of dietary sulfur nutrient input that is potentially linked to multiple deleterious effects on the mucosa, including the intestinal remodeling characterizing IBD and tumor risk. Bacterial production of sulfide through the metabolism of dietary and endogenous sulfur-containing compounds is ubiquitous in the gut microbiome^7,9,18,73,78^, but since it is not yet known which bacterial species have the greatest capacity for sulfide production, future studies will determine which species most contribute to altering gut sulfide levels.

### Dietary modification alters ISC function, including upregulation of sulfide detoxification pathways

We previously established that intestinal stem cells (ISCs) and lineages rapidly adapt to their nutritional environment by epigenetically altering pathways of energy and fat metabolism, as well as transport of calcium^28^. Detoxification of sulfide occurs through a multistep mitochondrial oxidation pathway, beginning with Sqrdl oxidizing sulfide^84,85^. We show that exposure to higher sulfide leads to changes in the mitochondrial sulfide oxidation pathway in Lgr5^hi^ ISCs within four days of shift to the NWD1 diet compared to mice fed AIN continuously. We observed increased expression of genes encoding the key enzymes Ethe1, Suox, and TST in the mitochondrial sulfide oxidation pathway in the mice fed NWD1, though the expression of Sqrdl, the transporter of sulfide into the mitochondria, was not different between the two diets. Since NWD1 alters Lgr5^hi^ stem cell gene expression, it will be important to determine the full spectrum of genes regulated through dietary alterations of the microbiome and sulfide production, and more broadly, how nonenzymatic gut redox chemistry can modify host gene expression and the luminal microenvironment as a whole.

Given the increased relative abundance of *Erysipelotrichaceae* in the mice, we next investigated whether this pattern was conserved in human patients with colorectal cancer. Because mouse and human microbiomes differ at the species level, we used the family level signal from the mice and applied it to the human metagenome dataset. We identified several *Erysipelotrichaceae* species that were associated with CRC, including the known CRC-associated bacteria *Solobacterium moorei*. Importantly, the presence of conserved early microbial changes in mice prior to tumor development and in humans with established tumors suggests that measurable microbial community perturbations present prior to the overt ecosystem disruption associated with the development of a neoplastic state.

### Dynamic/time series studies of diet/microbiome interactions to model environmental exposures and risk

This study focuses on temporal dynamics of dietary alterations in bacteria; the microbial metabolite, sulfide; and gut epithelial gene expression; as such, we acknowledge several limitations in our study. Though fed a diet that induces sporadic colon cancer in approximately 20% of mice, the mice in this study were not followed to the age at which they would develop intestinal tumors. Since age also affects microbiome composition^86^, the interaction between aging and dietary alterations in modifying microbiome composition may be complex. While we observe these alterations in mice, human stool sulfide is not routinely measured, nor are changes to the epithelium in response to elevated sulfide. With regards to the meta-analysis, the studies included patients from both Western and non-Western countries, so we cannot distinguish whether the association of *Erysipelotrichaceae* with CRC patients is a consequence of consuming of a Western diet, which causes a predisposition to CRC, or a factor directly influencing CRC development and/or progression. As such, *Erysipelotrichaceae* cannot be determined to have a causal role in CRC.

Our findings support a model in which environmental exposures such as diet reshape the gut ecosystem well before the development of an overtly neoplastic state. This includes measurable changes to microbiome composition, microbial metabolism, and epithelial gene expression. Notably, that microbial changes are present in the mice prior to the development of tumors indicates that ecosystem-level disruptions are measurable long before a pronounced disease state and may represent a potential point of intervention before a pathological ecosystem shift.

---

Author names in bold designate shared co-first authorship.

**Supplementary Figure 1:**
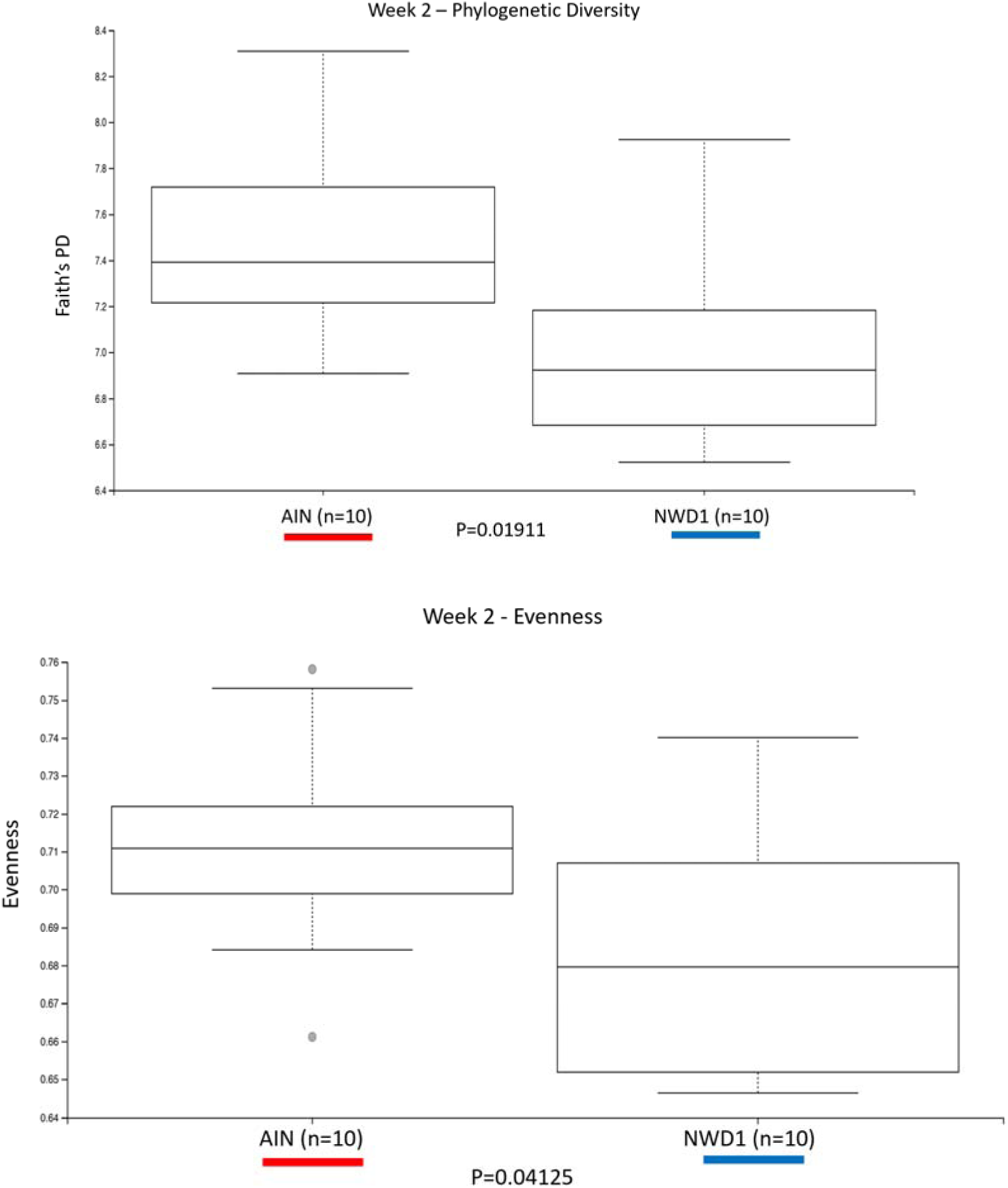
Week 2 Alpha Diversity & Evenness. Microbial communities from mice fed AIN show higher phylogenetic diversity (Kruskal-Wallis, p= 0.01911) and evenness than those fed NWD1 (Kruskal-Wallis, p=0.04125).

**Supplementary Figure 2:**
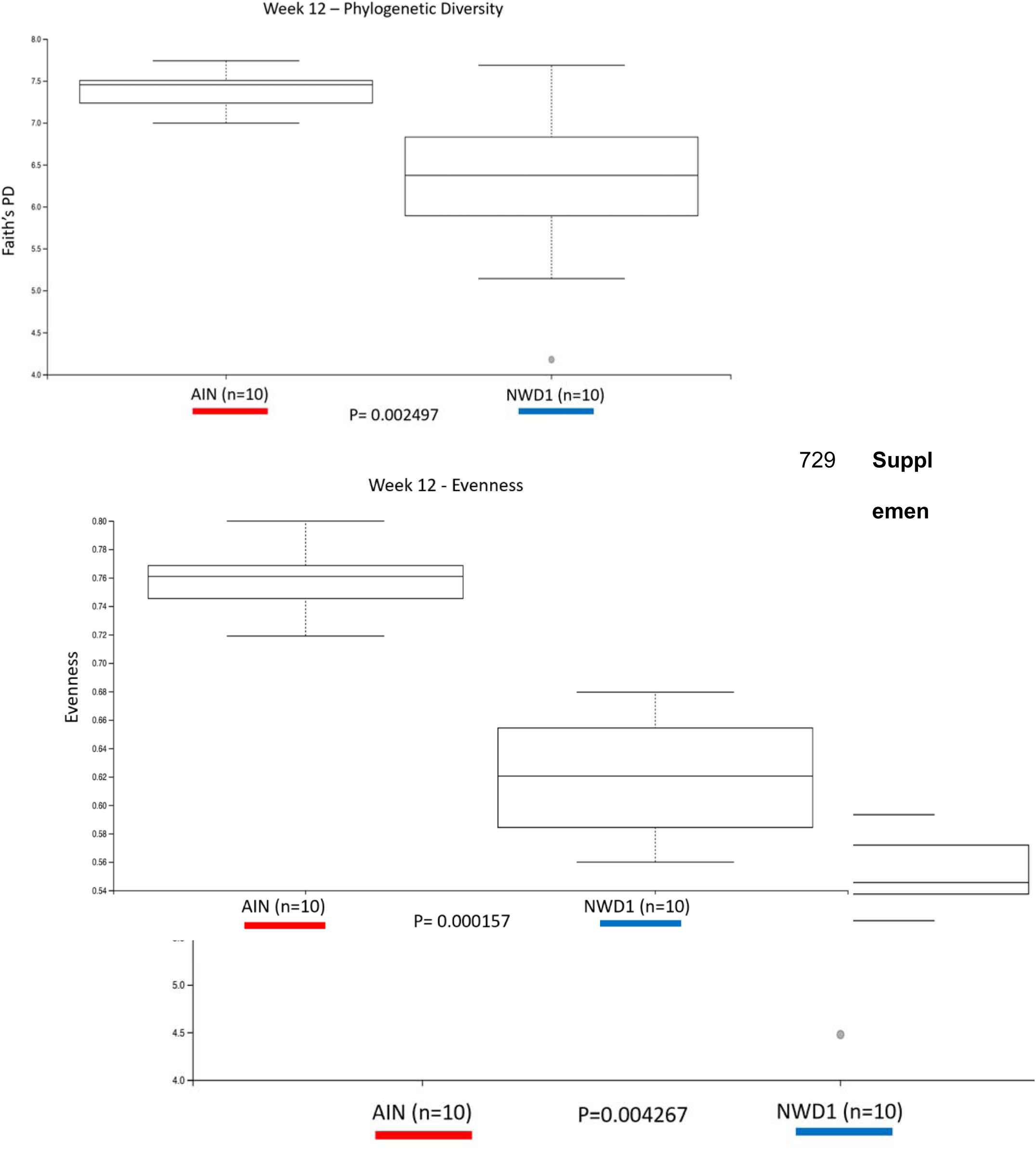
Week 12 Alpha Diversity & Evenness. Microbial communities from mice fed AIN show higher phylogenetic diversity (Kruskal-Wallis, p=0.002497) and evenness (Kruskal-Wallis, p=0.000157) than those fed NWD1.

**Supplementary Figure 3:**
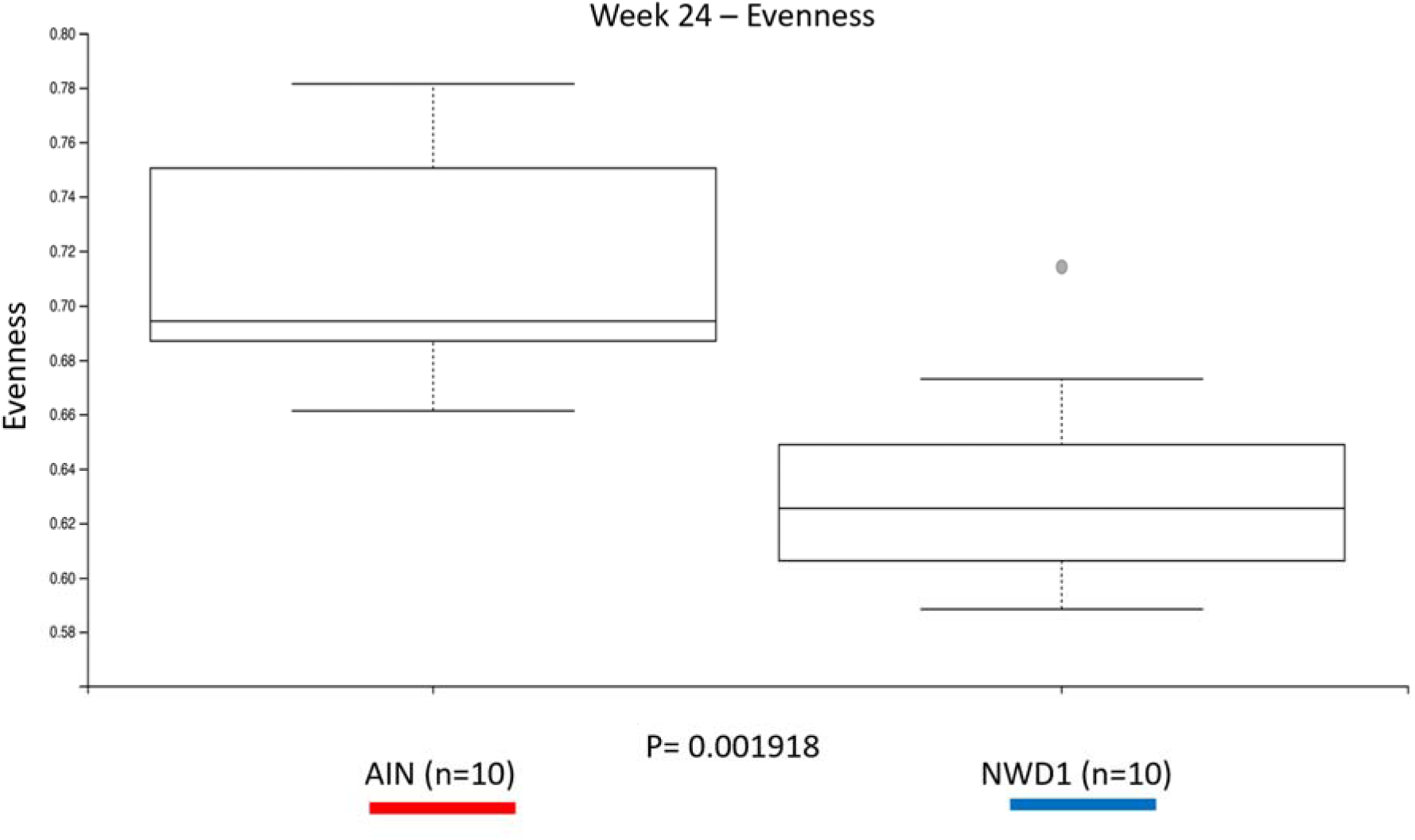
Week 24 Alpha Diversity & Evenness. Microbial communities from mice fed AIN show higher phylogenetic diversity (Kruskal-Wallis, p=0.004267) and evenness (Kruskal-Wallis, p= 0.001918) than those fed

**Supplementary Figure 4:**
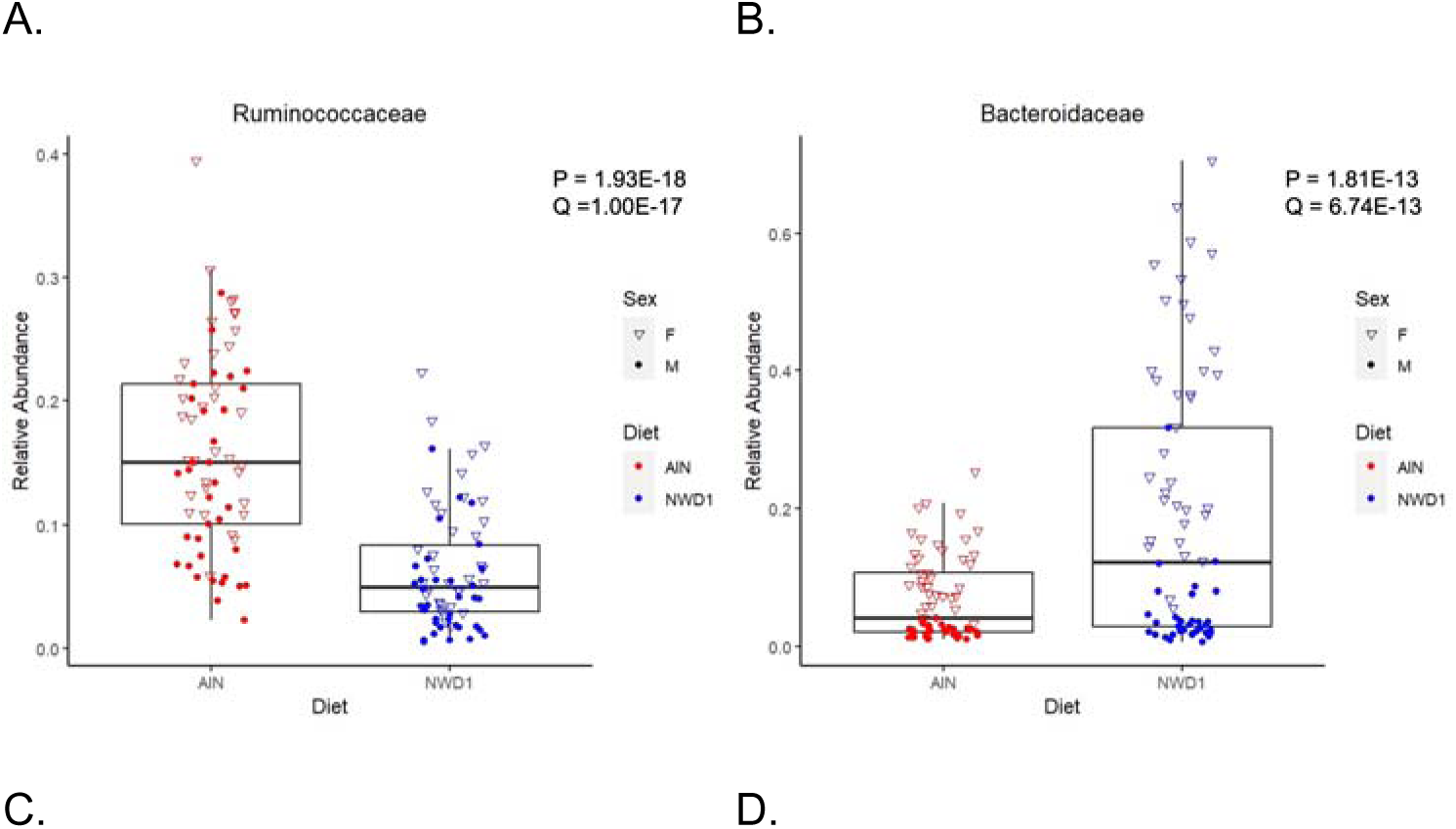
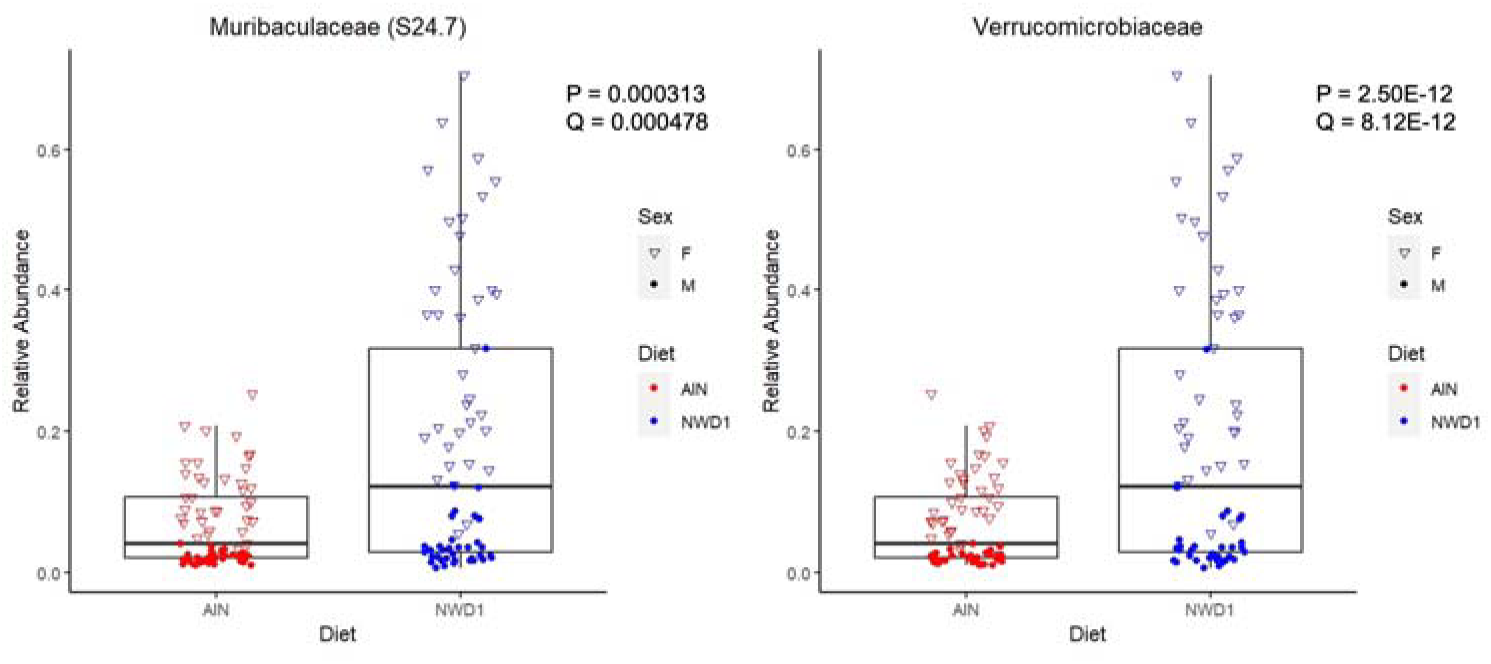
Relative abundances of the most abundant bacterial families in AIN- and NWD1-fed mice.

**Supplementary Figure 5.**
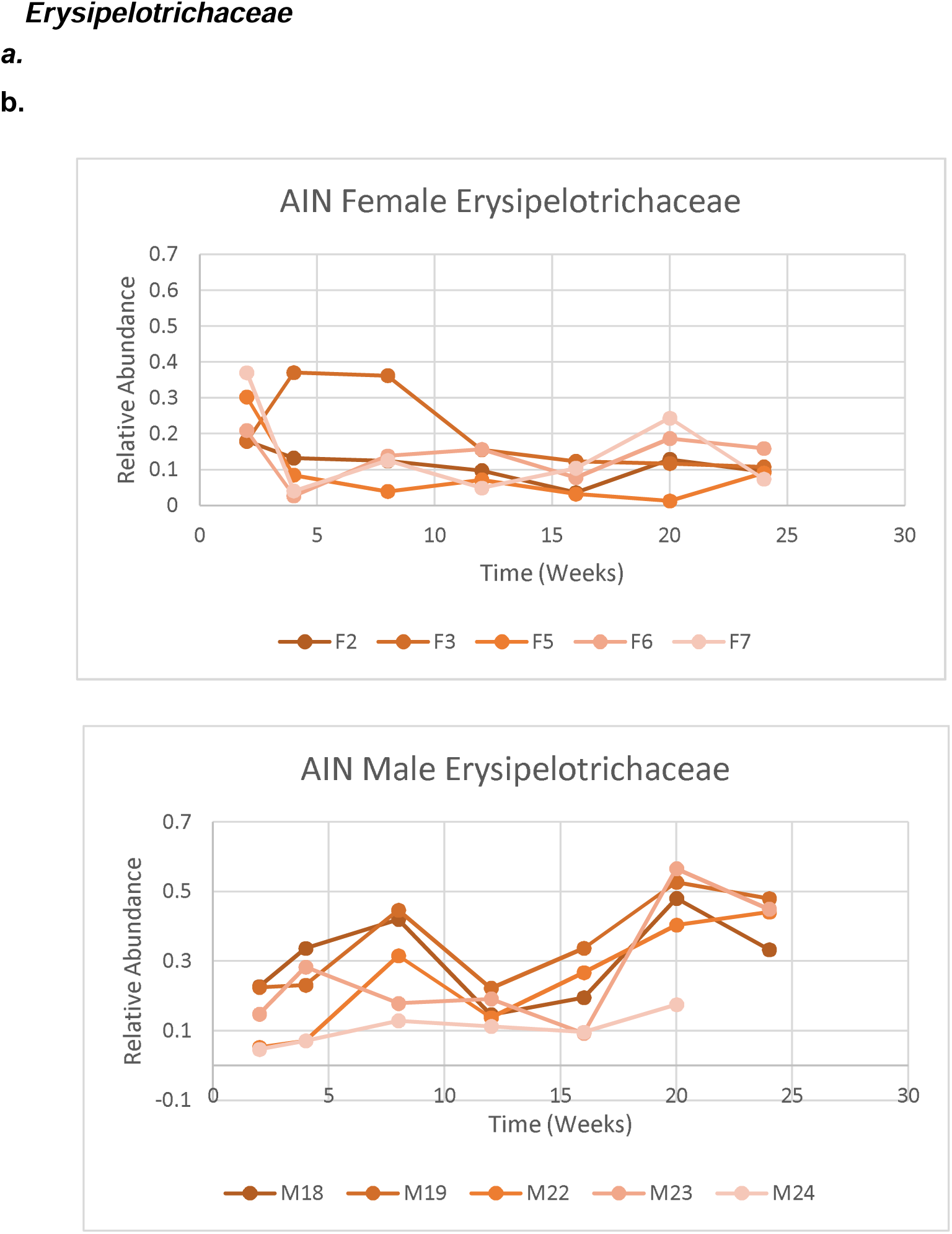
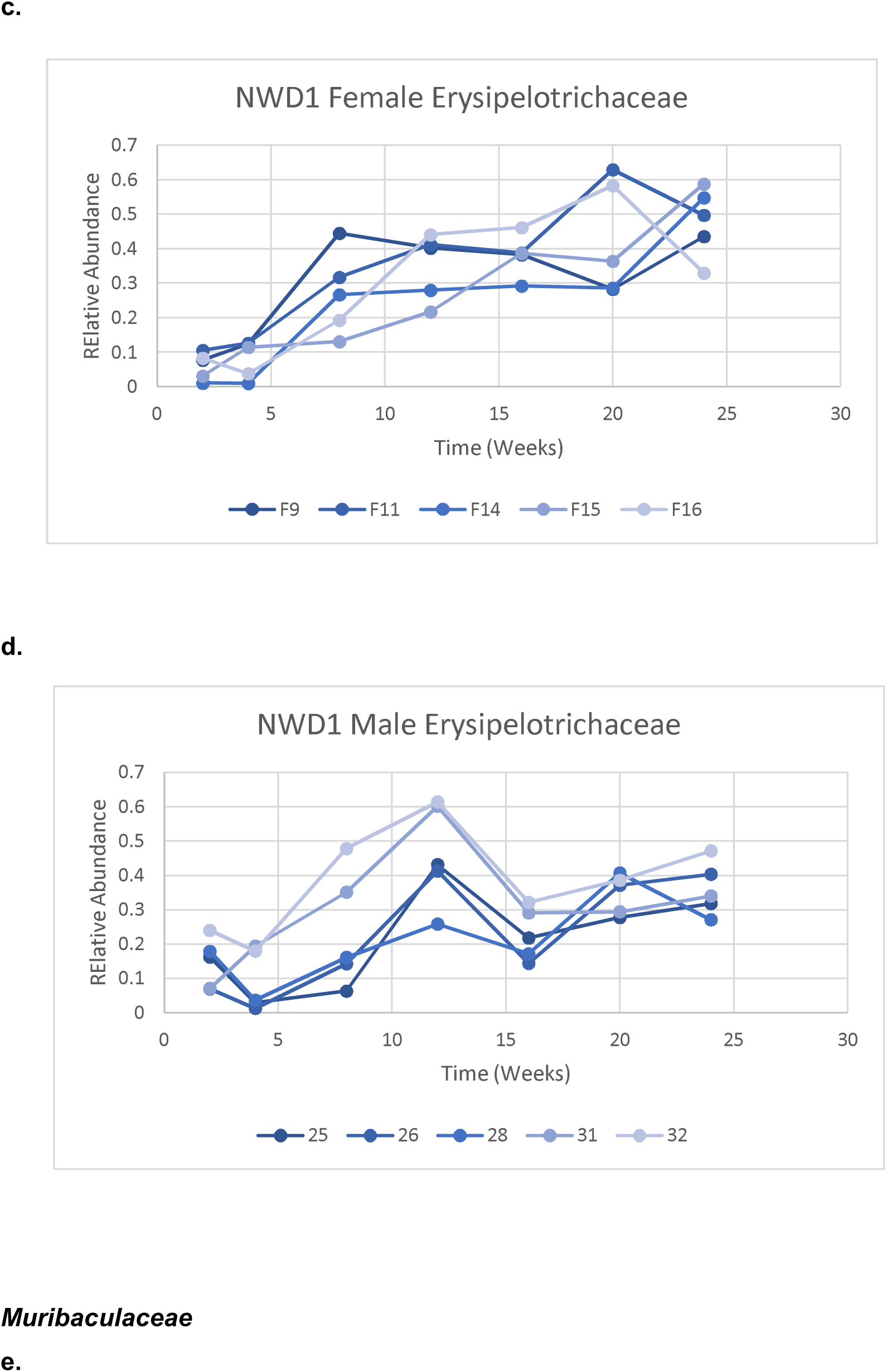
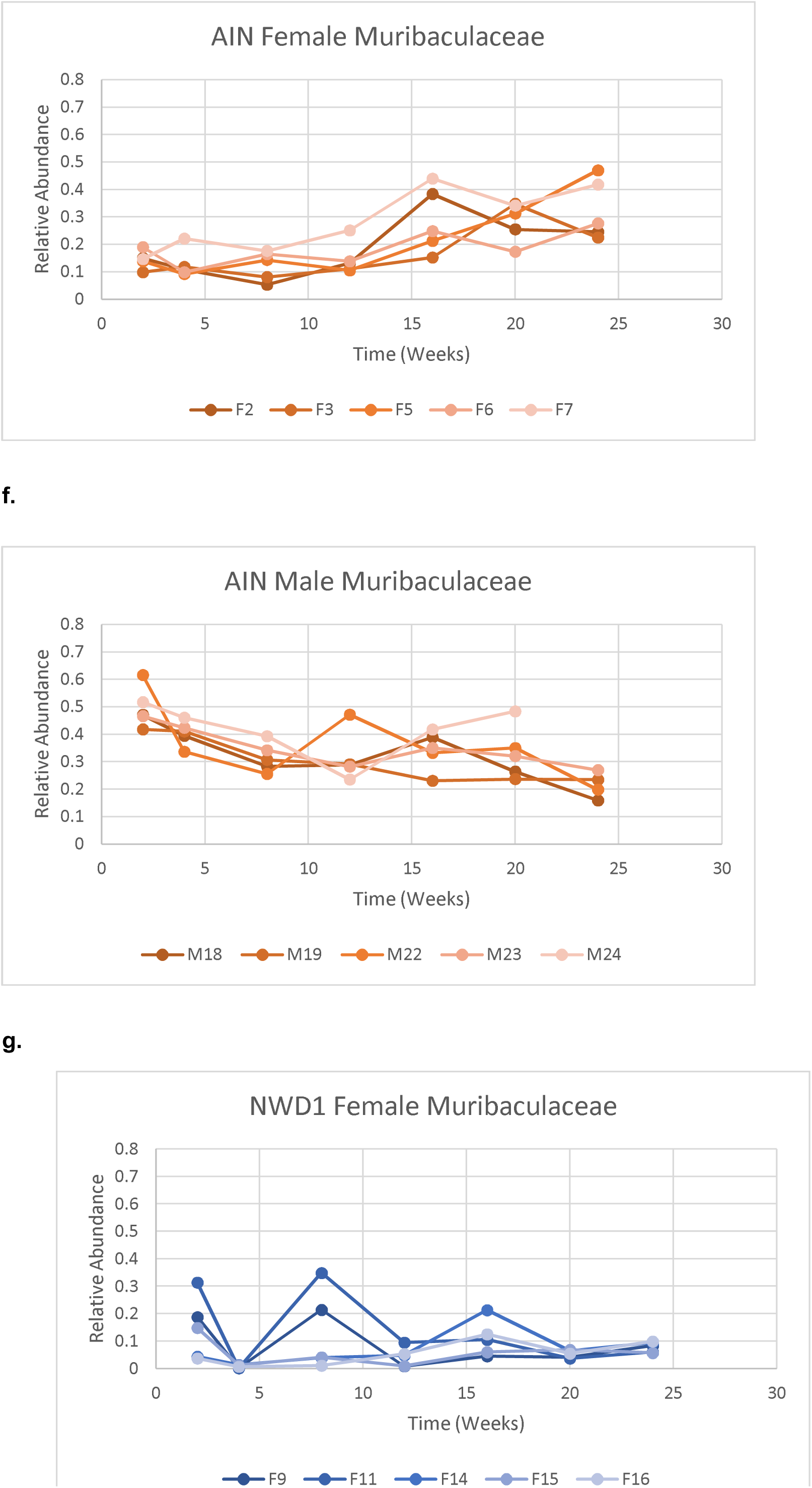
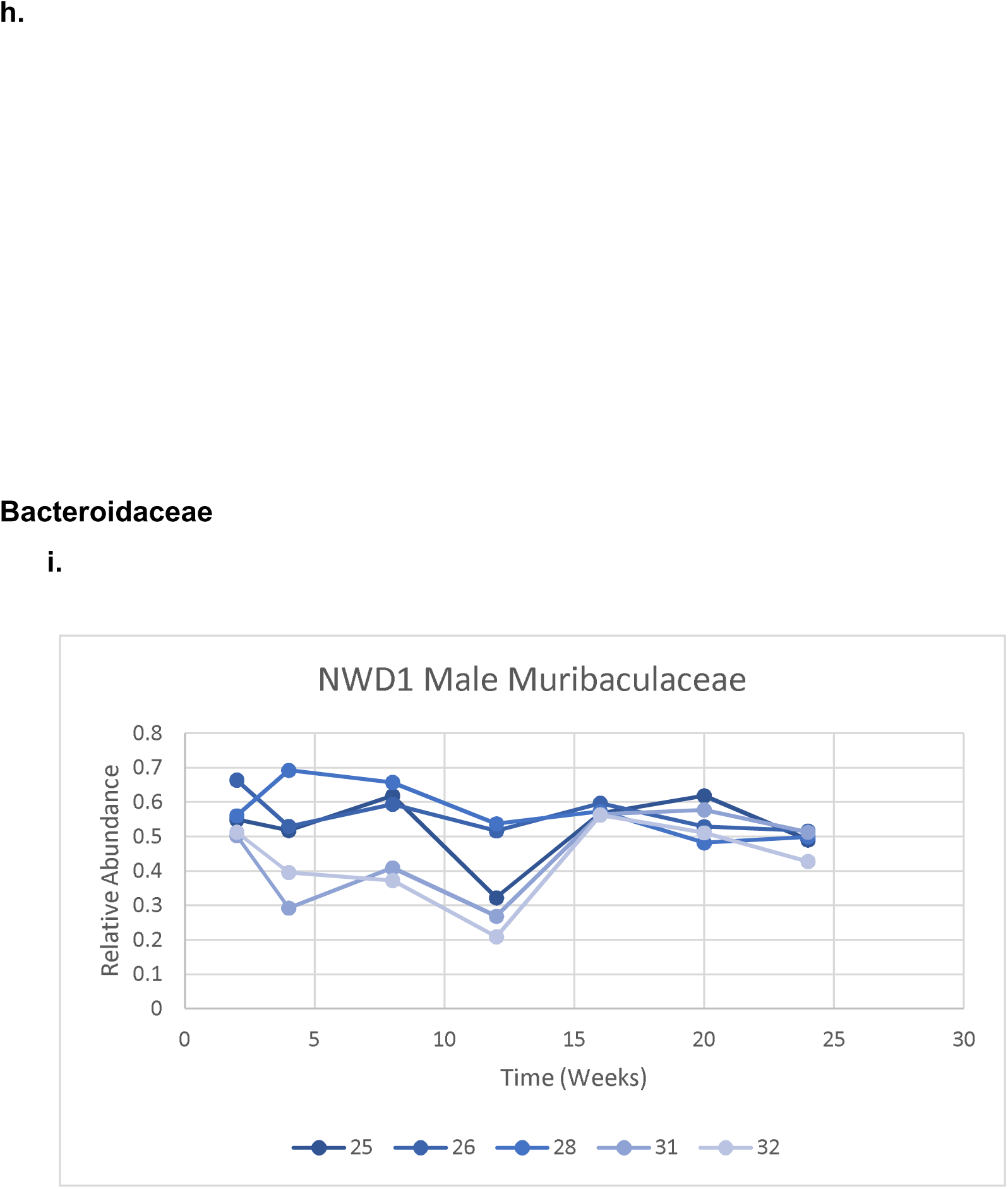
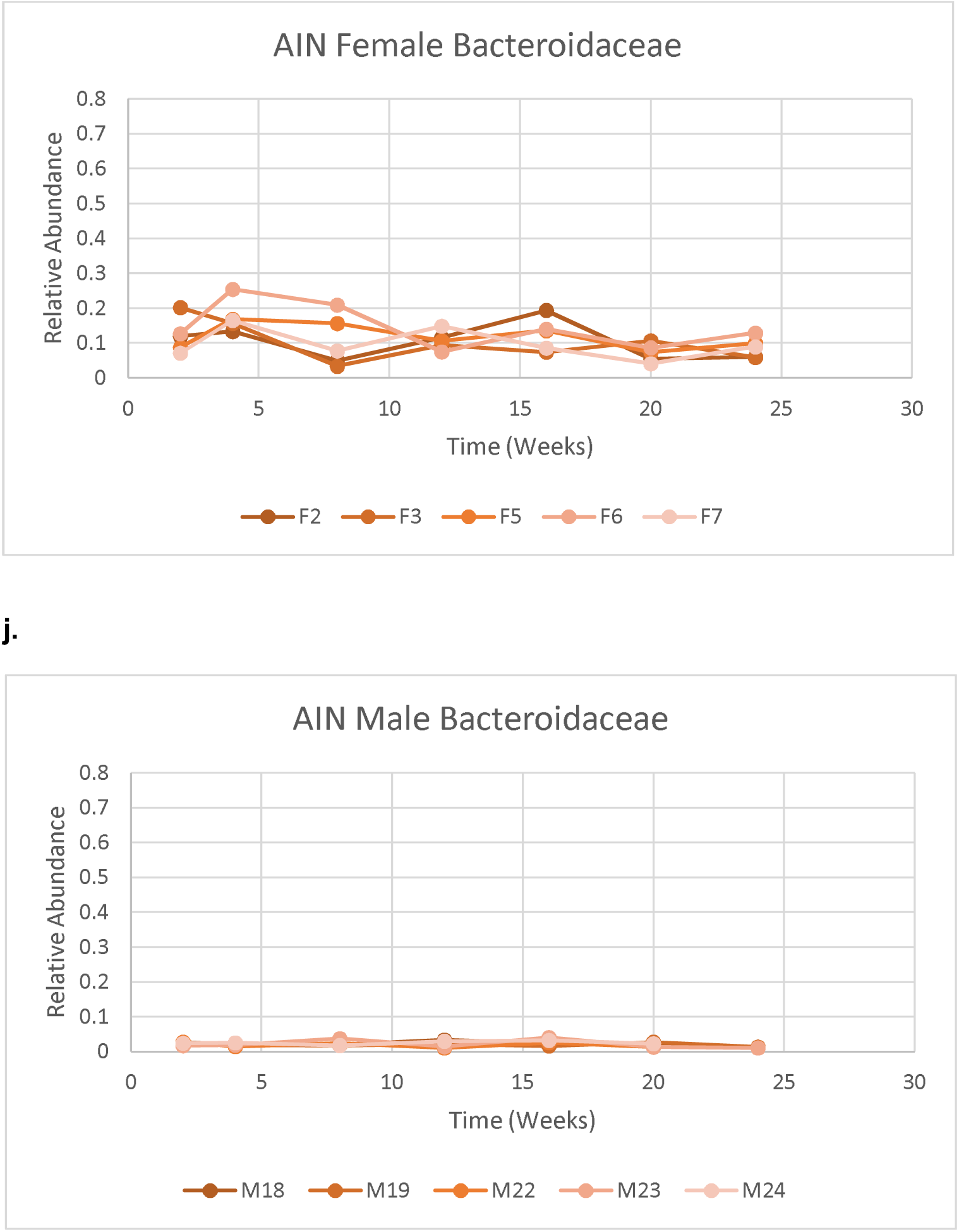
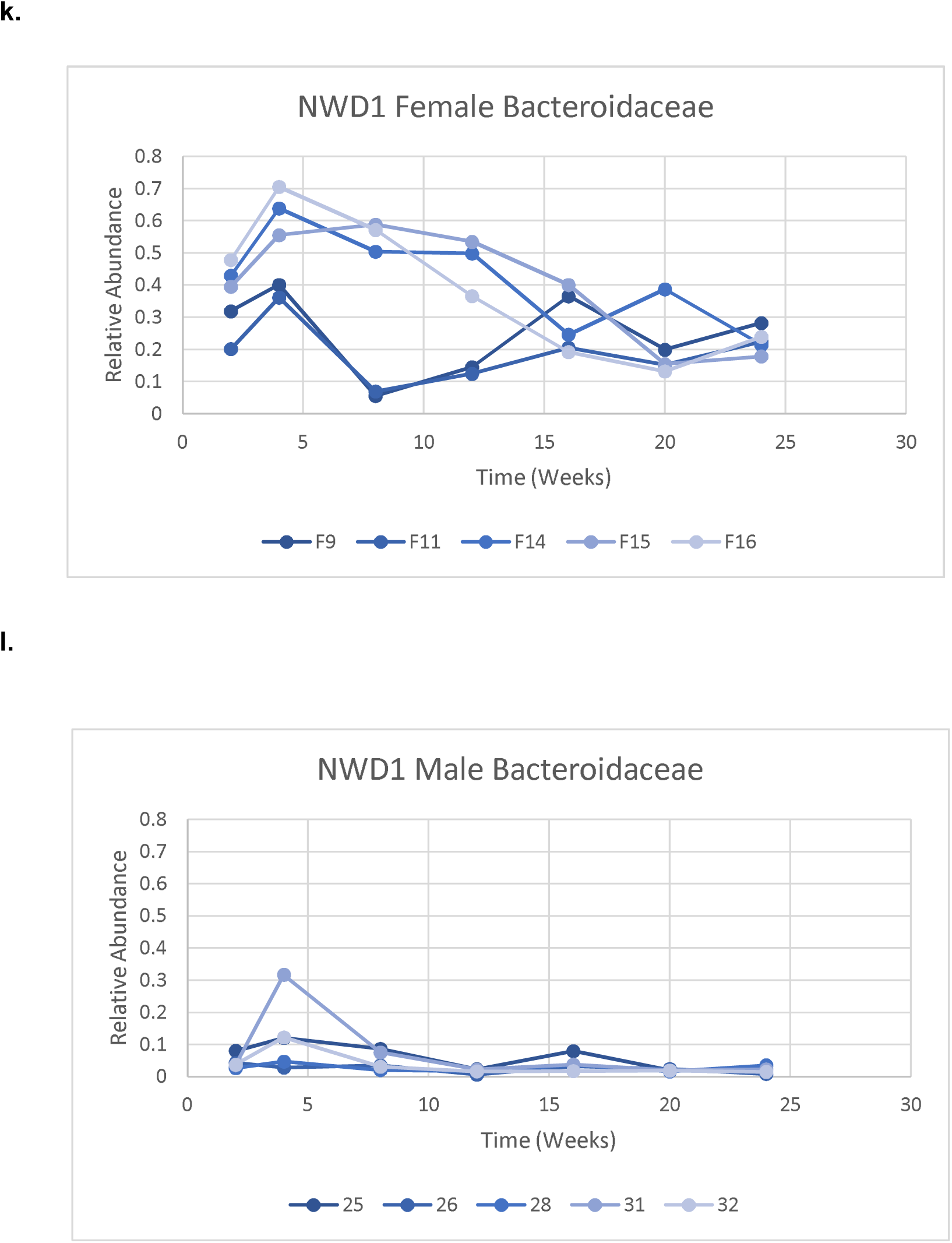
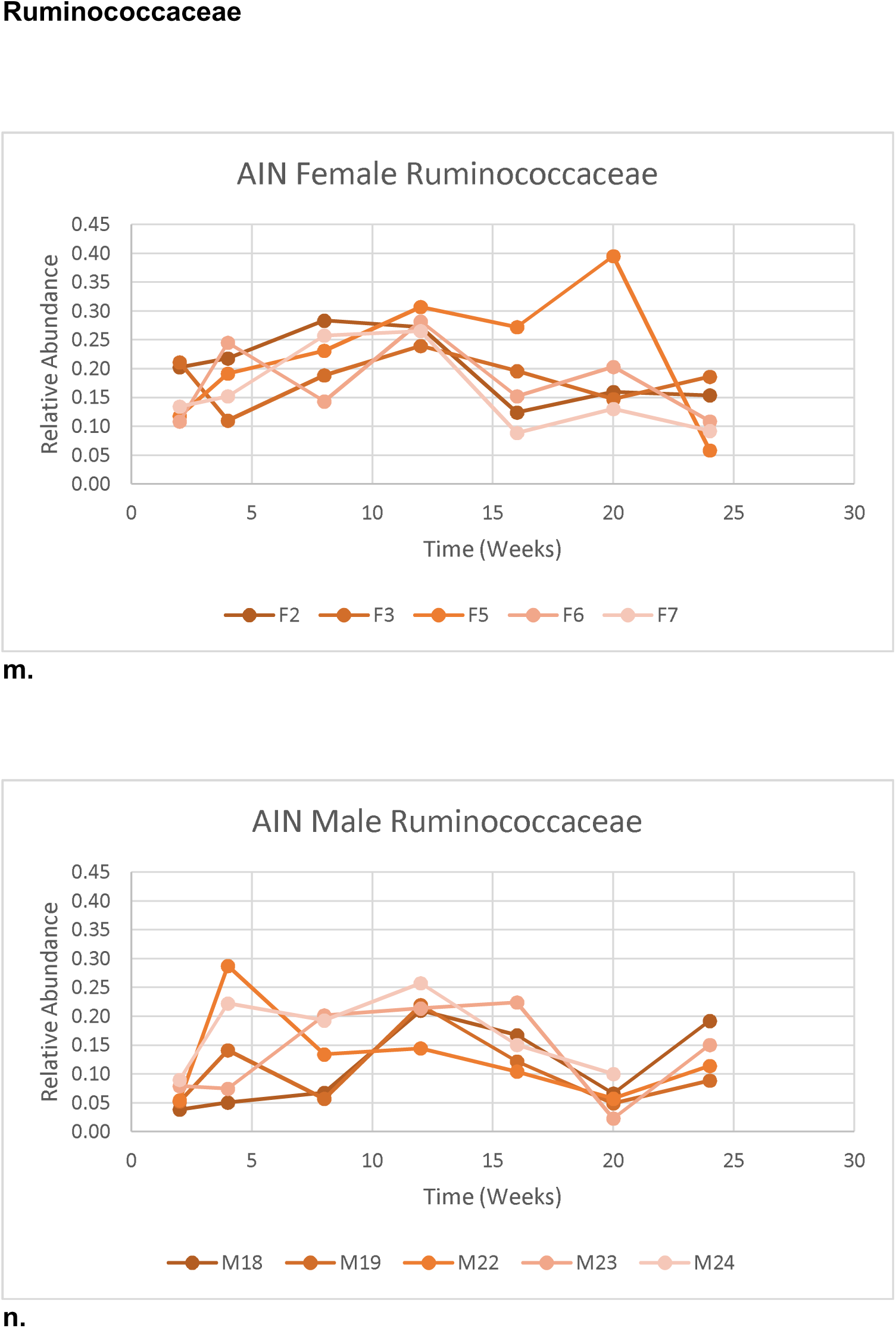
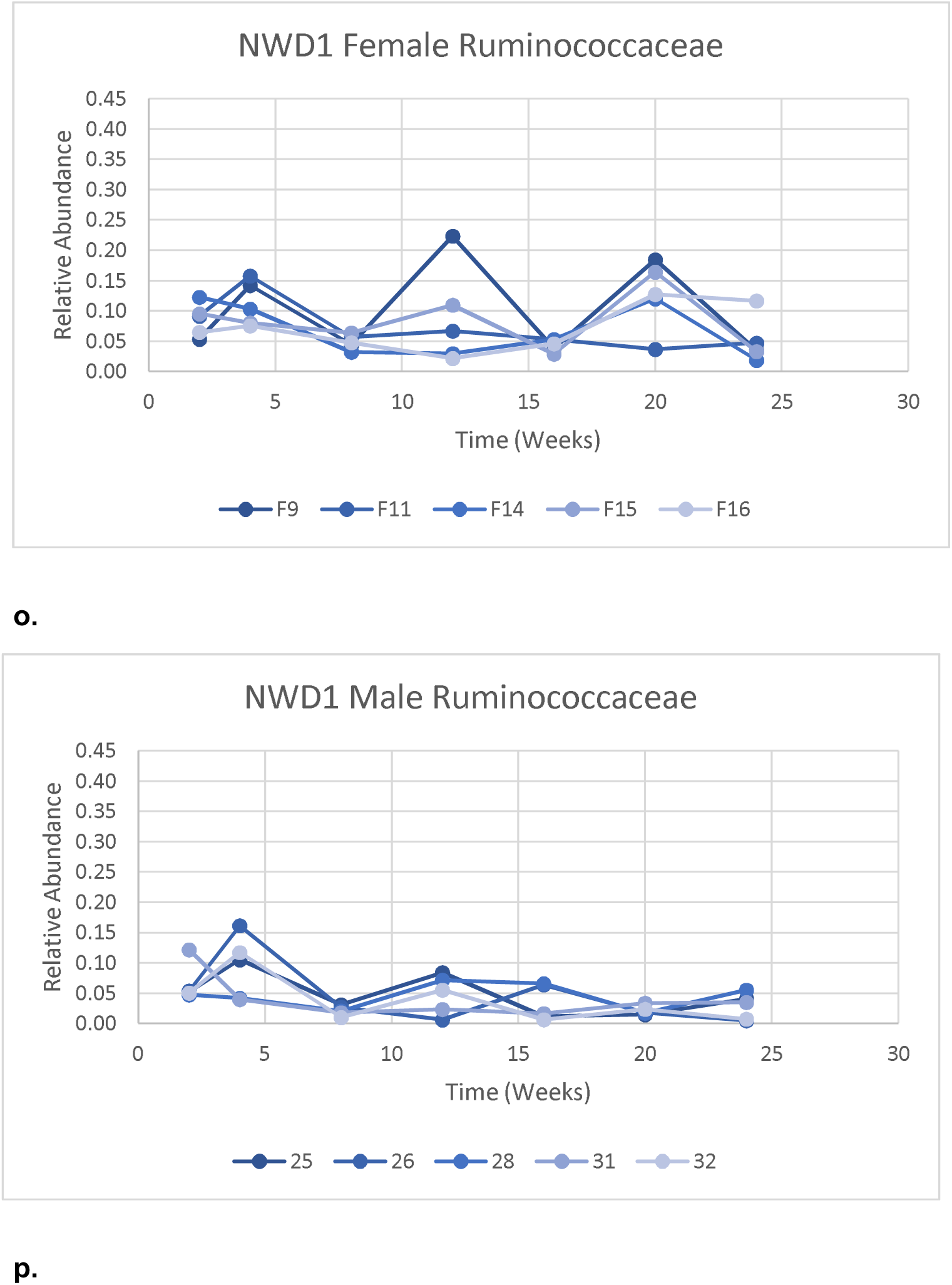
Relative Abundances of Bacterial Taxa Over Time in Mice Fed AIN or NWD1. Male (n=10) and female mice (n=10) were randomized to be fed AIN or NWD1 and their fecal microbiome composition was assessed longitudinally for 24 weeks. Relative abundances of the top four most abundant bacterial taxa were plotted to visualize changes over time in each individual mouse.

**Supplementary Table 1.**
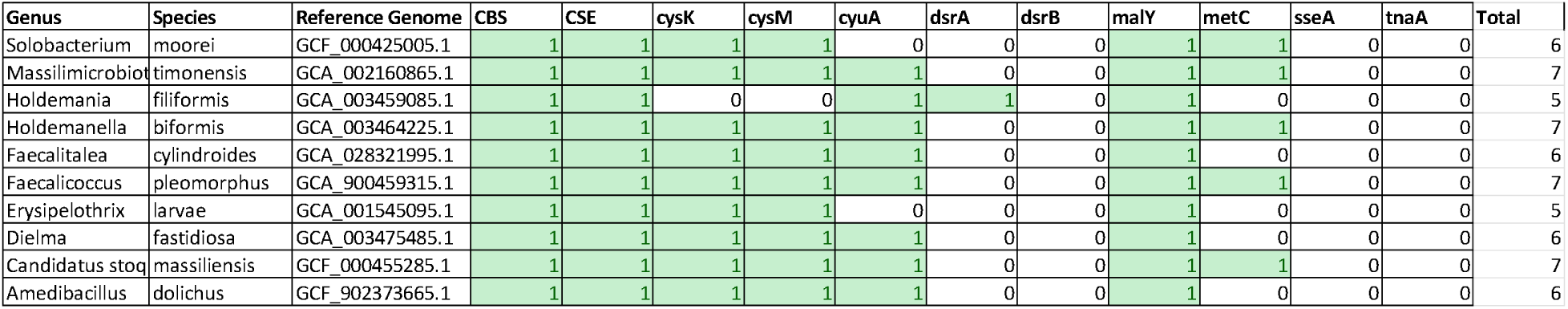
Sulfide-Producing Genes in *Erysipelotrichaceae*. To identify the presence or absence of sulfidogenic genes in *Erysipelotrichaceae* taxa, databases of homologous sequences for cysteine desulfhydrases and the dissimilatory sulfite reductase (dsrAB) genes were created and searched against each reference genome with a stringent cutoff of 1□×□10−10. Every species queried had between five and seven sulfidogenic genes, indicating the redundancy of this function.

## Supplementary Methods

### Meta-Analysis for Metagenomic Data Sets for CRC

Let *G* = {*g*_1_, *g*_2_,…, *g_J_*} be the set of studies included in our meta-analysis, where each *g_i_* represents a study. Each *g_i_* is itself a set of samples {*x^i^*_1_,*x^i^*_2_,…, *x^i^_m_*}, and each *x^i^_j_* is represented as a point in ℝ*^N^*, where *N* is the number of features used to annotate the sample (for example, *N* is the number of OTUs used to annotate a metagenomic sample). Additionally, each *x^i^_j_* may have some label *y^i^_j_* which describes which group the sample belongs in (case, control, etc.). We are interested in understanding how statistics computed on *g_j_* differ from study to study. Specifically, we want to identify features for which the statistics are consistently similar, in this case microbiome features that are more associated with CRC than with the healthy controls.

Let *S*: *Γ* → ℝ*^n^*, *Γ* ɛ *G* be a function which maps studies in *G* to real numbers and represents a statistical test. For example, *S* could be a function which returns the Z-score computed by comparing the values of samples in the study across groups. Denote *S^j^_n_* = *S_n_*(*g_j_*) be the value of the statistical test for the *n*-th feature when performed on study *j*.

With this outlined, our goal is to identify features *η* = {*n_l_*} such that for a given *n* ɛ *η*, all of the *S^j^_n_*,*j* ɛ [1, *J*] are “similar” in some way.

In Stouffer’s method, *S* is the function which computes a Z-score, and similarity is defined by asking if the mean of all the *S^j^_n_* is statistically different from 0. This captures the notion that, for the feature in question, the association between that feature and the target label is consistently in the same direction across studies. The association may actually be incomplete (meaning statistically below the significance threshold) in a given study, but we find that these incomplete associations tend to fall in the same direction, suggesting that they indicate a true association which we identify by this method.

In this paper, we take *S* to be the function which returns the AUC-score from a Mann-Whitney U test. This score represents the probability that a random sample taken from the control group will have a higher value than a random sample taken from the experiment group. This function is well-suited for microbiome data for a few reasons:

1. Microbiome samples are heavily zero-inflated. Many individuals simply do not carry a given organism at all. This by itself establishes that the data is not normally distributed.
2. Of the individuals that do carry an organism, their relative abundances are still not normally distributed. They are thought to be log-normal or inverse-binomial distributed because of how the sequencing works.

In principle, we could use a statistical test which attempts to model this distribution directly, such as a zero-inflated inverse-binomial. However, in Khan and Kelly 2023^87^, we show that even by using a less-biased (and hence weaker) test, we can identify a robust common signature of health and disease. The Mann-Whitney U-test is fully nonparametric, so it is safe to use in this setting without us having to verify any further distributional assumptions, at the cost of some statistical power. We make up this cost with the volume of data.

Additionally, the AUC-score is a single number (per feature) independent of the number of samples or the degrees of freedom in the study. This number is calculated from the actual U-score of the Mann-Whitney test as

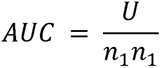

Where *n*_1_, *n*_2_ are the number of samples in the control and case groups. With this in mind, we can define similarity by asking for the average AUC score to be statistically different from 0.5 after making a normality assumption on the asymptotic distribution of the AUC score.

This method has been shown to be statistically efficient when the number of datasets is large and where datasets have similar variances. If some datasets have higher variance than others (for example, due to having less samples), they can inject noise into the final result, reducing the statistical power of this method. Liptak proposed a solution to this problem by weighting each study by some measure of the noise, for example, the square root of the study size, and this has empirically been shown to improve statistical power with these types of methods.

Hence, our final statistical test is as follows. We compute the AUC score for each microbiome feature across each CRC study *g_j_*, and we weight those scores by 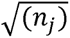. Let 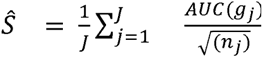. We then perform a one-sided t-test on *S* to reject the null hypothesis 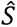 using a significance cutoff of α = .01. We denote those features for which the null hypothesis can be rejected at this threshold as general markers of health (when 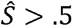) and disease (when 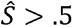).

## Abbreviations

AIN: AIN76A diet
CRC: colorectal cancer
Ethe1: ethylmalonic encephalopathy 1
H_2_S: hydrogen sulfide
IACUC: Institutional Animal Care and Use Committee
NWD1: New Western Diet 1
sqrdl: sulfide-quinone reductase-like protein
suox: sulfite oxidase
tst: thiosulfate transferase/rhodanese

## Author contributions

L.A. and L.K. conceptualized the study. Z.C., J.C., K.P., and C.S. collected data. Z.C., J.C., and S.W. analyzed the data. Z.C., L.A., and L.K. interpreted the data, drafted the manuscript, and revised the manuscript. All authors reviewed the manuscript.

